# Placentrex disrupts the biofilm formation of *Pseudomonas aeruginosa* through multi-target transcriptional reprogramming

**DOI:** 10.64898/2026.05.22.727083

**Authors:** Bayomi Biju, Ajit R Sawant, Sachin Maji, Tejavath Ajith, Tridib Neogi, Piyali Datta Chakraborty, Anindya Sundar Ghosh

**Affiliations:** Department of Bioscience and Biotechnology, Indian Institute of Technology Kharagpur, Kharagpur, West Bengal-721302, India; Department of Research and Development, Albert David Limited, 5/11, D. Gupta Lane. Kolkata, West Bengal-700050, India

**Keywords:** Pseudomonas aeruginosa, Placentrex, biofilm, preformed biofilm, EPS

## Abstract

**Aims:** *Pseudomonas aeruginosa* biofilm-associated infections pose a significant clinical challenge due to their inherent antibiotic tolerance. This study aimed to evaluate the antibacterial and antibiofilm activity of Placentrex, a standardised aqueous placental extract, against *P. aeruginosa* and to elucidate its molecular mechanism of action using RNA sequencing (RNA-seq).

**Methods and Results:** Placentrex exhibited potent bactericidal activity against P. aeruginosa at 50 µg/mL. Biofilm formation was significantly inhibited by ∼87% at 50 µg/mL after 72 hours. Preformed biofilms were eradicated by ∼93% and ∼89% at 50 and 25 µg/mL, respectively. Interestingly, biofilm viability was reduced by ∼93% and ∼87% upon treatment with 50 µg/mL and 25 µg/mL of Placentrex, respectively. EPS characterisation revealed that the EPS contain a single large polysaccharide, and chromatography data suggested that it is made up of glucose as a monomer. RNA-seq identified coordinated downregulation of seven key genes, namely, flp major pilin (surface attachment), extracellular solute binding protein (ABC transporter-mediated nutrient sensing and biofilm maintenance), gntP permease (carbon metabolism), AraC family transcriptional regulator (quorum sensing and polysaccharide biosynthesis), ureE (urease nickel metallochaperone), aromatic amino acid permease (pyoverdine and PQS biosynthesis), and MFS transporter (efflux and autoinducer export).

**Conclusions:** Placentrex exerts comprehensive antibiofilm and antibacterial activity through simultaneous disruption of surface attachment, nutrient-sensing-driven biofilm maintenance, quorum sensing, carbon metabolism, urease virulence maturation, and efflux-mediated persistence. This polypharmacological mechanism supports Placentrex as a promising multi-target antibacterial agent against P. aeruginosa biofilm-associated infections.

**Impact statement:** Placentrex is a potential anti-biofilm agent against Pseudomonas aeruginosa.

## Introduction

Healthcare-associated infections, especially those caused by multidrug-resistant Gram-negative (MRGN), represent an increasingly critical challenge in infection control, contributing significantly to morbidity and mortality while offering only limited therapeutic options (Vonberg et al., 2008; Wellington et al., 2013). One such MRGN, which is the leading cause of nosocomial infections posing a severe threat to immunocompromised individuals, including those with cystic fibrosis, burns, chronic obstructive pulmonary disease (COPD), or cancer, as well as patients reliant on ventilators or invasive medical devices (Thi et al., 2020), is *Pseudomonas aeruginosa*. *Pseudomonas aeruginosa* is a Gram-negative, opportunistic pathogen known for its metabolic versatility, environmental ubiquity, and clinical significance. Additionally, its ability to form biofilms enhances resistance to antibiotics and host immune responses, and also contributes to persistent and recurrent infections. *P. aeruginosa* employs a vast arsenal of virulence factors, including quorum sensing, type III secretion systems (T3SS), elastases, pyocyanin, and rhamnolipids, which facilitate tissue invasion, immune evasion, and cytotoxicity (Qin et al., 2022). The adaptive defence mechanism, like efflux pumps, β-lactamase production, and horizontal gene transfer, along with the genomic flexibility, categorise it as a multi-drug resistant (MDR) ESKAPE pathogen by WHO. The rise of carbapenem-resistant and extensively drug-resistant (XDR) strains of *P. aeruginosa* has rendered many conventional therapies ineffective, exacerbating mortality rates in healthcare settings (Mendes Pedro et al., 2024). These resistance mechanisms, referred to as the three lines of defence by Zou et al., are classified based on their site of action. The primary or outermost line comprises bacterial biofilm, followed by cell wall, cell membrane, and housed efflux pumps, and finally the intracellular environment, where biochemical and genetic responses play a critical role in conferring resistance once all preceding barriers have been breached (Zhou et al., 2015).

Biofilms are microbial communities that form a typical protective spatial architecture, adhere to one another and to the surfaces surrounding them, and develop relationships with the host, adapt to hostile external conditions, and show resistance to antibiotics and other environmental cues (Rather et al., 2021). The biofilm architecture is sustained by a self-synthesised extracellular polymeric substance (EPS) that includes polysaccharides, matrix proteins, and extracellular DNA (eDNA) (S. Wang et al., 2015). Biofilm formation is a multifaceted cyclic process that progresses through five stages. The process is initiated when a stressed planktonic cell makes a reversible contact with a surface through the flagellum or cell pole, followed by irreversible attachment marked by reduced flagellar gene expression and matrix deposition (Sauer et al., 2022). This initial colonisation is influenced by surface roughness, cell surface hydrophobicity and covalent-ionic interactions between the cell and surface. This is followed by the maturation stage, where more cells are recruited into the matrix-embedded cluster to multiply and form microcolonies and three-dimensional structures with nutrient-delivering channels. In the final stage, cells disperse either actively in response to stress such as biofilm aging, nutrient limitation or antimicrobial exposure, or passively due to external forces like fluid shear or mechanical disturbance (Zhao et al., 2023). This stage is important for bacterial survival, spread, and disease transmission.

Unlike the free-floating planktonic bacteria, bacteria inhabiting the protected environment of a biofilm differ in character. For instance, biofilm bacteria confer up to 1,000-fold greater resistance to antibiotics compared to planktonic bacteria. This resistance not only leads to chronic and recurrent infections but also contributes significantly to the global rise in antimicrobial resistance, making conventional therapies increasingly ineffective(Roy et al., 2017; Shariati et al., 2022; Srinivasan et al., 2021). As a result, there is a critical demand for innovative agents that can effectively disrupt biofilms and enhance the efficacy of existing antimicrobial treatments.

Placentrex, a human placental extract, has gained attention for its multifaceted therapeutic properties, particularly in wound healing, immunomodulation, and as an adjunct in infection management. Its use as an antimicrobial agent is increasingly justified due to the limitations of conventional antibiotics, especially in the context of chronic infections, where biofilm formation by pathogens significantly impedes treatment efficacy(Nishan, 2020; Sharma, 2016). Placentrex is a pharmacological preparation derived from human placental tissue. It contains a complex blend of bioactive compounds, including peptides (such as fibronectin type III and ubiquitin-like moieties), nucleotides, amino acids, enzymes, vitamins, minerals, growth factors, and cytokines (Chakraborty et al., 2012) that contribute to its therapeutic effects. Notably, studies have demonstrated that placental peptides possess direct antimicrobial activity, inhibiting the growth of various microbes in vitro as well as biofilm inhibition comparable to established agents like chlorhexidine, particularly against *Enterococcus faecalis* and *Escherichia coli*(Bandyopadhyay et al., 2018). The integration of Placentrex into standard wound care protocols could represent a significant advancement in the management of antibiotic-resistant, biofilm-mediated infections. Thus, placentrex represents a promising agent in the fight against biofilm-associated infections, offering a multifactorial approach that addresses both microbial eradication and biofilm management, which traditional antibiotics cannot. Therefore, understanding the antibiofilm and antimicrobial potential of Placentrex, particularly against *Pseudomonas aeruginosa*, is essential to advance strategies for combating biofilm-mediated resistance and to evaluate its potential as an effective adjunct in antimicrobial therapy.

## Materials and methods

### Bacterial strains, culture media, and chemicals

The bacterial strain used in this study was *Pseudomonas aeruginosa* MTCC 424 (PA 424). The bacterial strain was grown at 37 °C in Luria–Bertani broth (LB) or LB agar (Hi-Media, Mumbai, MH, India). Cation-adjusted Mueller–Hinton Broth (MH) was used for all experiments (HiMedia, Mumbai, MH, India). Human placental extract, Placentrex, was provided by Albert David Ltd, Kolkata, India. Overall, the manufacturing procedure holding confidentiality of the proprietary terms has been described earlier.(Chakraborty et al., 2012). Unless otherwise specified, all the other reagents, including antibiotics, were purchased from Sigma-Aldrich (St. Louis, MO, USA).

### Determination of susceptibility towards Placentrex

Antimicrobial susceptibility of PA to Placentrex was determined according to the Clinical and Laboratory Standards Institute (CLSI) guidelines, using the microbroth dilution method (Lewis II et al., 1986).

### Time□kill assay

The viable cell count method was used for the time-kill assay. PA cell suspension of OD_600nm_ ∼0.2 was incubated in the presence of different concentrations of Placentrex. Aliquots were removed at 0, 24, 48, and 72 hours post-inoculation and serially diluted in 0.85% sodium chloride solution for the determination of viable counts. 100μL of the diluted samples were plated in duplicate on LB agar plates and incubated at 37°C for 18 hours. The time taken to kill the initial bacterial load was assessed by plotting the log CFU/mL versus incubation time (Petersen et al., 2006).

### Semi- quantitative crystal violet-based biofilm inhibition assay

The anti-biofilm activity of Placentrex against PA was assessed using the crystal violet staining method. Different concentrations (50 µg/mL- 1.56 µg/mL) of Placentrex were introduced in the wells of a sterile 24-well polystyrene microtitre plate containing the media with OD_600nm_ ∼0.2 bacterial inoculum, incubated at 37°C for 72 hr under static conditions. Following this, the wells were washed three times with phosphate buffer saline (PBS) to remove the planktonic cells and stained with 0.1% crystal violet (CV) stain for 15 min. Following staining, the excess CV was removed by washing with water. The bound dye was solubilised in 1 ml of 33% acetic acid (v/v), and the absorbance of the acetic acid-dye solution was measured at 600 nm using an ultraviolet/visible spectrophotometer [Multiskan Spectrum-1500 Spectrophotometer (Thermo Scientific, Switzerland)] (Pal et al., 2019). The percentage inhibition in biofilm formation was calculated by the formula [(C-T)/C]*100, where T is the stain released from the test wells, and C is the stained control wells(Wu et al., 2019).

### Qualitative analysis of biofilm inhibition

a. **Bright-field microscopy** To assess the biofilm production, a 24 well plate polystyrene plates fitted with microscopic coupons containing different concentrations (50 µg/mL- 1.56 µg/mL) of Placentrex and media were inoculated with OD_600nm_ ∼0.2 bacteria at 37°C for 72 hr under static conditions The biofilm grown on coupons was washed thrice with 1X PBS (phosphate-buffered saline), dried and stained with 0.1%crystal violet for 15 min and were washed with water. The biofilms formed on the coverslips were subsequently visualised using an OLYMPUS 1 ×71 (Olympus Corporation, Tokyo, Japan) (Pal et al., 2019).
b. **Field emission scanning electron microscopy** Biofilm was grown as before on microscopic coupons, fixed with 2% glutaraldehyde, dried, and mounted on aluminium stubs. The samples were shadowed with gold and analysed through a ZEISS EVO 60 scanning electron microscope (Pal et al., 2019).

### Semi- quantitative crystal violet-based preformed biofilm eradication assay

To understand the effect of Placentrex on the preformed biofilms, OD_600nm_ ∼0.2 bacterial inoculum was introduced in the wells of a sterile 24-well polystyrene microtitre plate containing the media, incubated at 37°C for 72 hr under static conditions for biofilm formation. After the incubation, the spent media were discarded, and the wells were washed thrice with 1X PBS. Fresh media and Placentrex at different concentrations were added to the wells and incubated for 24 hours (G. Kumar et al., 2019). The spent media was discarded the next day, and the wells were washed with autoclaved water and kept for drying. The wells were dried at room temperature and stained with 0.1% crystal violet, and the biofilm was quantified as mentioned previously (Pal et al., 2019; Wu et al., 2019).

### Biofilm viability assay

The viability of bacterial cells within the biofilm was determined using the dye triphenyl tetrazolium chloride (TTC). The preformed biofilms were treated with Placentrex as above and incubated for 24 hours. After incubation, the wells were washed, and fresh culture media (MH) were added and incubated for 6-7 hours at 37 °C. Further, the viability of cells was analysed by incubating 200µL of this culture with 0.05% TTC at 37°C for 15 minutes in the dark, and the change in colour was measured using Multiskan GO microplate spectrophotometer at 490nm (Thermo Fisher Scientific) (G. Kumar et al., 2019).

### Fluorescent microscopic analysis of biofilm viability

For fluorescence microscopy, the biofilm was developed on coupons and treated with different concentrations of placentrex as in the preformed biofilm eradication assay. The coupons were stained with BacLight Live/Dead dye as per the manufacturer’s protocol (Invitrogen Inc., CA, USA) and visualised using OLYMPUS IX71 (Olympus Corporation, Tokyo, Japan)(Kumar et al. 2012, 2015).

### Twitching and swimming motility assays

The twitching motility of PA was assessed on a semisolid medium containing 10g/L pancreatic digest of casein, 5g/L sodium chloride, 6g/L beef extract, and 0.8% agar. 20 mL of medium containing different concentrations of Placentrex (50 µg/mL- 12.5 µg/mL) and TTC dye (MacFaddin, 2000) was poured into a sterile petri dish, bacterial culture was stabbed on the surface of the medium, and incubated at 37°C for 24h. The procedure of the swimming motility assay was consistent with the twitching motility assay except that the swimming medium, containing 0.4% agar and bacterial culture, was stabbed at the interface between the bottom of the petri dish and the medium(Kelly & Fulton, 1953).

### Microbial Adhesion to Hydrocarbons (MATH) assay

Bacterial cell surface hydrophobicity will be assessed using ‘microbial adhesion to hydrocarbons’ (MATH). Under the conditions used, the hexadecane–water interface represents a hydrophobic substratum surface, with a net negative charge due to adsorption of negative ions from the aqueous phase. Briefly, overnight-grown PA cells were inoculated in media containing Placentrex (50 µg/mL- 12.5 µg/mL). After 5 hours of incubation, cells were washed and resuspended to an OD at 600 nm (*A*_0_) between 0.4 and 0.6 in PUM buffer (100 mM K2HPO4, 50 mM KH2PO4, 30 mM urea, 1 mM MgSO4). To the suspension, 500μl of hexadecane was added. The suspensions were vortexed for 3-5 minutes and left to stand at room temperature to allow the layers to separate. Absorbance of the aqueous layer was measured at 600nm. The hydrophobicity index (HPBI) or % hydrophobicity was calculated as the change in percentage from the original absorbance (Ao) to the final absorbance (Af) upon hexadecane exposure [% hydrophobicity = (A_0_ − Af)/A_0_ × 100](A. Kumar et al., 2015; Van Der Mei et al., 2003).

### Isolation of EPS from culture broth

EPS was isolated as described by Bales et al. (2013) with minor modifications. Briefly, an OD_600nm_ ∼0.2 bacterial inoculum was incubated in 25 mL of medium containing different concentrations of Placentrex (50 µg/mL- 12.5 µg/mL). Biofilms were grown at 37°C without shaking for 4–5 days to obtain a thick slurry of biofilm. After the development of a mature biofilm, 150 ml of formaldehyde (36.5% solution) was added to the sludge to fix the cells and to prevent the cells from lysing during subsequent steps. This mixture was incubated at room temperature with gentle shaking (100 rpm) for 1 hour. Ten mL of 1 M NaOH was added to the same mixture, and incubated at room temperature with shaking for 3 hours to extract EPS. After incubation, the suspension was centrifuged (16,8006g) for 1 hour at 4°C. The supernatant containing EPS was filtered through a 0.2 mm filter and dialysed against distilled water using a 12–14 kDa molecular weight cut-off (MWCO) membrane for 24 hours at 25°C. Following dialysis, 20% w/v TCA was added to the extracted EPS solutions on ice for 30 minutes to remove most of the protein and nucleic acids. The solution was centrifuged at 16,800g for 1 hour at 4°C, and the supernatant was collected. To the supernatant, 1.5 volumes of 95% ethanol were added, and incubated at -20 °C for 24 hours to precipitate exopolysaccharides away from lipids. The solution was then centrifuged at 16,800g for 1 hour at 4°C and the exopolysaccharide pellet was resuspended in Milli-Q water and dialysed against the same for 24 hours at 4°C using a 12–14 kDa MWCO membrane to remove low-molecular-weight impurities. The remaining retentate was lyophilised and stored at-20°C until further use (Bales et al., 2013).

### Determination of total sugar in EPS by phenol-sulphuric acid assay

The total sugar content of the lyophilised EPS samples was estimated using a phenol-sulfuric acid assay (Dubois et al., 1956) by comparing it with a standard glucose curve. For the assay, 20μl of 1mg/mL of lyophilised EPS in 0.1M sodium hydroxide solution was mixed with 200μl of concentrated H_2_SO_4_ and 20μl of 5 % phenol in a 96-well microtiter plate and mixed gently on a dancing shaker for 5-10 minutes. The colour change in samples was determined spectrophotometrically at *A*490nm (Multiskan GO microplate spectrophotometer; Thermo Fisher Scientific). Concentrations of total sugar in EPS samples were calculated from the standard glucose graph.

### Determination of total protein by Bradford assay

The total protein in the EPS samples was estimated by Bradford assay(Bradford, 1976) with slight modifications. Briefly, 200μl of 5mg/mL of lyophilised EPS in 1X PBS was mixed with 2 mL of commercially available Bradford reagent, and incubated in the dark for 10 minutes at room temperature. After incubation, the absorbance of the samples was determined spectrophotometrically at *A*595nm. The total protein concentration in EPS samples was calculated from the standard BSA graph.

### Determination of the average molecular weight of different polysaccharides in the EPS samples

The approximate average molecular weight of the exopolysaccharide was determined using the gel filtration chromatography method(Hara et al., 1982). To determine the molecular weight of EPS, a Sepharose CL-6B column was used. Before loading, the column was stabilised in 0.1 M sodium hydroxide. The isolated EPS samples (1mL of 10mg/mL in 0.1 M sodium hydroxide) were loaded onto the column and eluted at room temperature by using 0.1 M sodium hydroxide solution as the mobile phase. Eluate was collected in 5 ml fractions, totalling one column volume, and the presence of polysaccharides was subsequently determined using the phenol-sulfuric acid assay (Dubois et al., 1956). Column calibration was performed using dextran standards with molecular weights ranging from 6 to 2000 kDa (Mondal et al., 2023). To visualise the presence of polysaccharides in each fraction, thin-layer chromatography was performed using butanol: acetic acid: water (4:2:2) as mobile phase and orcinol sulphuric acid reagent was sprayed and kept at 60 °C for 10–15 min to develop colour (Finocchiaro, 1966).

### Analysis of monosaccharides by thin-layer chromatography (TLC) and High-Performance Liquid Chromatography (HPLC)

All the fractions containing polysaccharides were pooled and dialysed against distilled water to remove residual NaOH and subsequently lyophilised to yield pure polysaccharides. The pooled fractions were estimated by the phenol sulphuric acid method to determine the polysaccharide concentration. These EPS samples were then hydrolysed using 1 ml of 4 M trifluoroacetic acid (TFA) at 120 °C for 4–5 h in an oil bath. Subsequently, the hydrolysates were neutralised by repetitive sample washing with methanol in a rotary evaporator at 50 °C until pH 7 was attained (Blakeney et al., 1983). These TFA-digested EPS samples were analysed for their monomeric sugar composition by TLC on a silica plate (TLC silica gel 60 F254; Merck) and compared with standard sugars. Here, acetone, butanol and water in the ratio of 5:4:1 were used as the mobile phase for the separation of the constituents. After the sample had run, the silica plate was air-dried and sprayed with orcinol-sulphuric acid solution as before (Finocchiaro, 1966).

The monosaccharide composition of the hydrolysed EPS samples was further confirmed and quantified by HPLC (Debnath et al., 2024), with minor modifications. After neutralisation, the samples were dissolved in Milli-Q water and analysed for monosaccharides by high-performance liquid chromatography (HPLC). For analysis of hydrolysates and monosaccharide standards by HPLC, Milli-Q water was used as the mobile phase; the column (Hiplex-H) temperature was 65°C, the RID temperature was set at 35°C, and the flow rate was 0.5 ml/min. Individual monosaccharides were quantified by comparing the peak regions to standard calibration curves made for each standard monosaccharide.

### RNA sequencing

The RNA was extracted from three biological replicates of the control (Untreated) and Placentrex-treated (12.5µg/mL) PA groups. The RNA-seq analysis was performed at Chromosome Labs Private Limited, Mumbai, India. The original raw data from the Illumina platform is transformed into sequenced reads by base calling. Raw data are recorded in a FASTQ file, which (Andrews, 2010)h contains sequence information (reads) and corresponding sequencing quality information. The quality of the reads was assessed using FastQC v0.11.9 (Andrews, 2010) before proceeding with downstream analysis. Raw reads were passed through fastp v0.23.2 (Chen, 2023) to remove adapters, low-quality reads (Q<30), and reads with length < 50 nucleotides after adapter trimming. Rockhopper at https://cs.wellesley.edu/~btjaden/Rockhopper/ was used for de novo transcriptome assembly, and assembly completeness was assessed by CheckM v1.2.4 (Parks et al., 2015). Bakta v1.11.4(Schwengers et al., 2021) was used to annotate transcripts using its database v6. The Reads were mapped against the assembly using Bowtie2 (v2.5.4) (Langmead & Salzberg, 2012). After mapping, quantification was performed with featureCounts v1.5.3 (Liao et al., 2014), and gene abundance estimation was performed with featureCounts v2.0.3. The total feature counts were 9881. Differential gene expression was analysed after batch correction using the DESeq function from the DESeq2(Love et al., 2014) package with a false discovery rate (FDR) cut-off of ≤ 0.05 and a minimum expression log2-fold change (FC) of ≥ ±2. The significantly differentially expressed genes were obtained from the Bakta-annotation and analysed using topGO (Alexa & Rahnenführer, 2021), with Pseudomonas Gene Ontology annotations used as the reference.

### Statistical Analysis

All the experiments were conducted thrice with three replicates in each set of experiments, and the results were calculated as mean ±standard deviation. The statistical significance (*P* value) was calculated using GraphPad by performing two two-sample, unpaired Student’s *t-test* where ns*P >0.05*, _∗_*P* ≤*0.05*, _∗∗_*P* ≤*0.001*, _∗∗∗_*P* ≤*0.0001*, and _∗∗∗∗_*P* ≤*0.0001*.

## Results

### Placentrex inhibits the growth of PA

The antibacterial activity of Placentrex is analysed by the microbroth dilution method. Placentrex showed growth-inhibitory effects against PA, with a minimum inhibitory concentration (MIC) of 50µg/mL.

### Placentrex instantaneously kills PA

The bactericidal effect of Placentrex on the planktonic growth of PA was investigated by determining the minimum bactericidal concentration (MBC), which was found to be 50 µg/mL. The effect of Placentrex against PA was investigated by studying time-kill kinetics (Fig.S1), and Placentrex showed concentration- and time-dependent bacterial killing ability. Exposure to Placentrex at 50 µg/mL (MIC) resulted in complete elimination of PA cells by 24 hours, and the effect was sustained through 72 hours. At 25 µg/mL (½ × MIC), complete eradication of viable bacteria was achieved by 48 hours and similarly persisted until 72 hours. Complete eradication of viable bacteria indicates irreversible cellular damage (Stojowska-Swędrzyńska et al., 2023), and the absence of bacterial regrowth throughout the observation period suggests that Placentrex may impede the development of resistance (Baquero & Levin, 2021).

### Placentrex reduced the biofilm-forming ability of PA

Inhibition of biofilm formation can be due to direct bactericidal effects or by inhibiting the attachment of viable bacterial cells to the surface. To evaluate this, the antibiofilm activity of Placentrex against PA was determined by growing the biofilms in the presence of different concentrations of Placentrex. We observed a concentration-dependent inhibition of biofilm formation after 72 hours of incubation. The biofilm forming was reduced by approximately ∼87%, ∼72%, and ∼58% in the presence of Placentrex at 50 µg/mL, 25 µg/mL, and 12.5 µg/mL, respectively (Fig. 1a and 1b). Ciprofloxacin (0.0625µg/mL) was used as a positive control. Inhibition of PA biofilms at sub-MIC concentration suggests that biofilm inhibition can be due to the inhibition in attachment to the surface, and that at higher concentrations can be due to the bactericidal effect.

**Figure 1.**
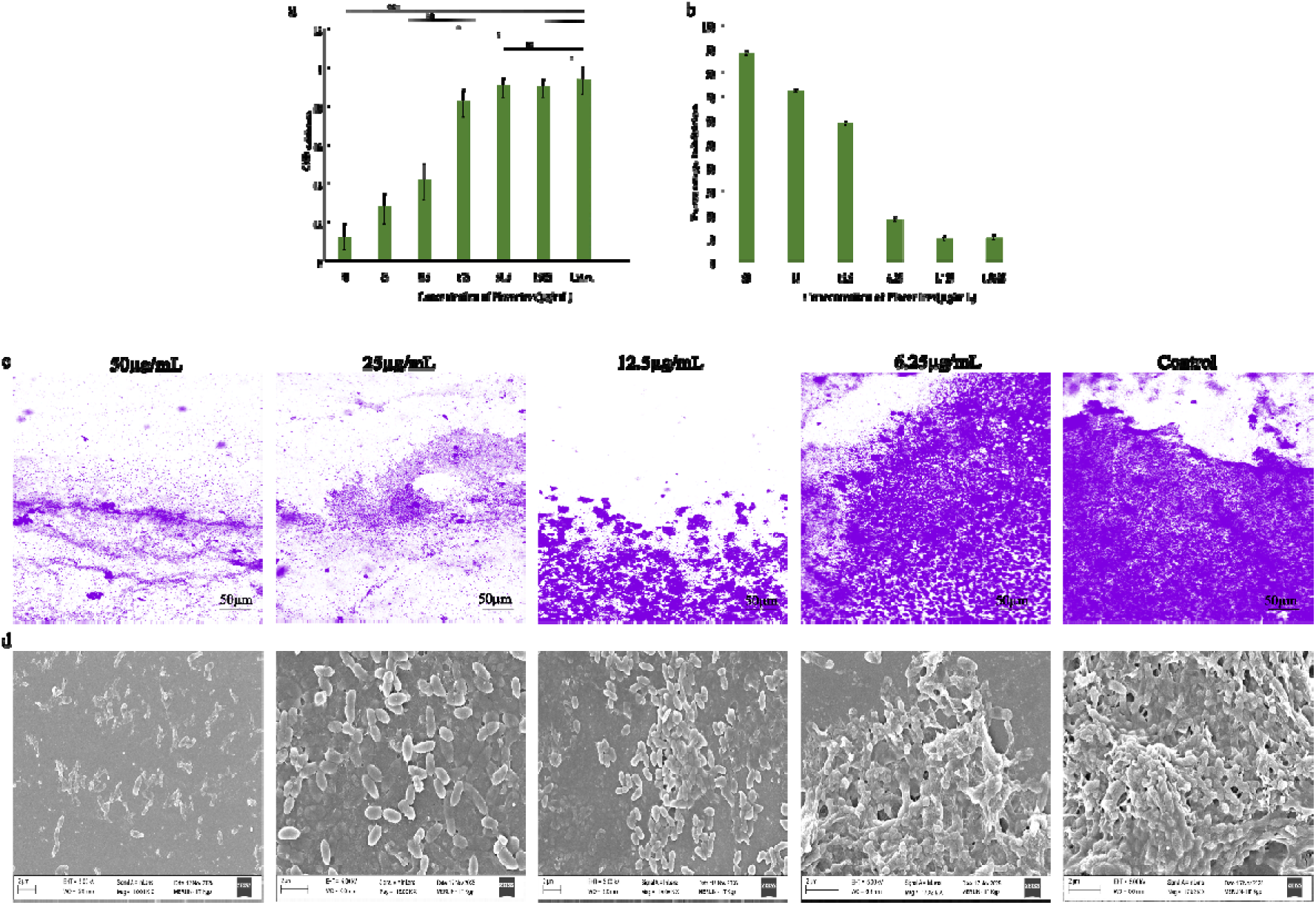
Semi-quantitative biofilm formation assay of PA cells. a) Biofilm formation by PA cells, b) Percentage inhibition of biofilm formation by PA cells in the presence of different concentrations of Placentrex by the Crystal Violet staining method. c) Bright field microscopy, d) field emission scanning electron microscopy images of biofilms formed by PA in the presence of different concentrations of Placentrex.

Further bright-field microscopic analysis substantiated the concentration-dependent biofilm reduction compared to the untreated biofilm. A distinct difference in biofilm-forming pattern and reduced biofilm density was observed with increasing concentration of Placentrex (Fig. 1c).

SEM analysis further confirmed the concentration-dependent inhibition of biofilm formation. The untreated control showed a dense, multi-layered biofilm with an extensive extracellular matrix embedding tightly clustered bacterial cells. In contrast, biofilms treated with Placentrex at 12.5 µg/mL exhibited a partial reduction in cell density with disruption of the matrix architecture. At 25 µg/mL, a markedly sparse biofilm was observed with isolated cell clusters and considerably diminished EPS. Treatment at 50 µg/mL resulted in severely compromised biofilm integrity, with individual cells showing morphological alterations suggestive of membrane damage, consistent with bactericidal activity at higher concentrations (Fig. 1d).

### Placentrex reduced the preformed biofilms and also reduced the viable bacteria within the biofilm of PA after 24 hours of treatment

Unlike planktonic cells, biofilms can withstand conventional antibiotics due to the presence of extracellular polymeric substances (EPS). Based on this, the ability of Placentrex to disperse preformed biofilm of PA was evaluated using the minimum biofilm eradication concentration (MBEC). According to the crystal violet assay, the MBEC of Placentrex was measured to be 12.5 µg/mL. Further, the preformed biofilm was cleared by ∼93% and ∼89% upon treatment with 50 mg/ mL and 25 µg/mL of Placentrex, respectively (Fig. 2a). Therefore, it can be inferred that Placentrex can effectively eradicate the preformed biofilm of PA.

**Figure 2.**
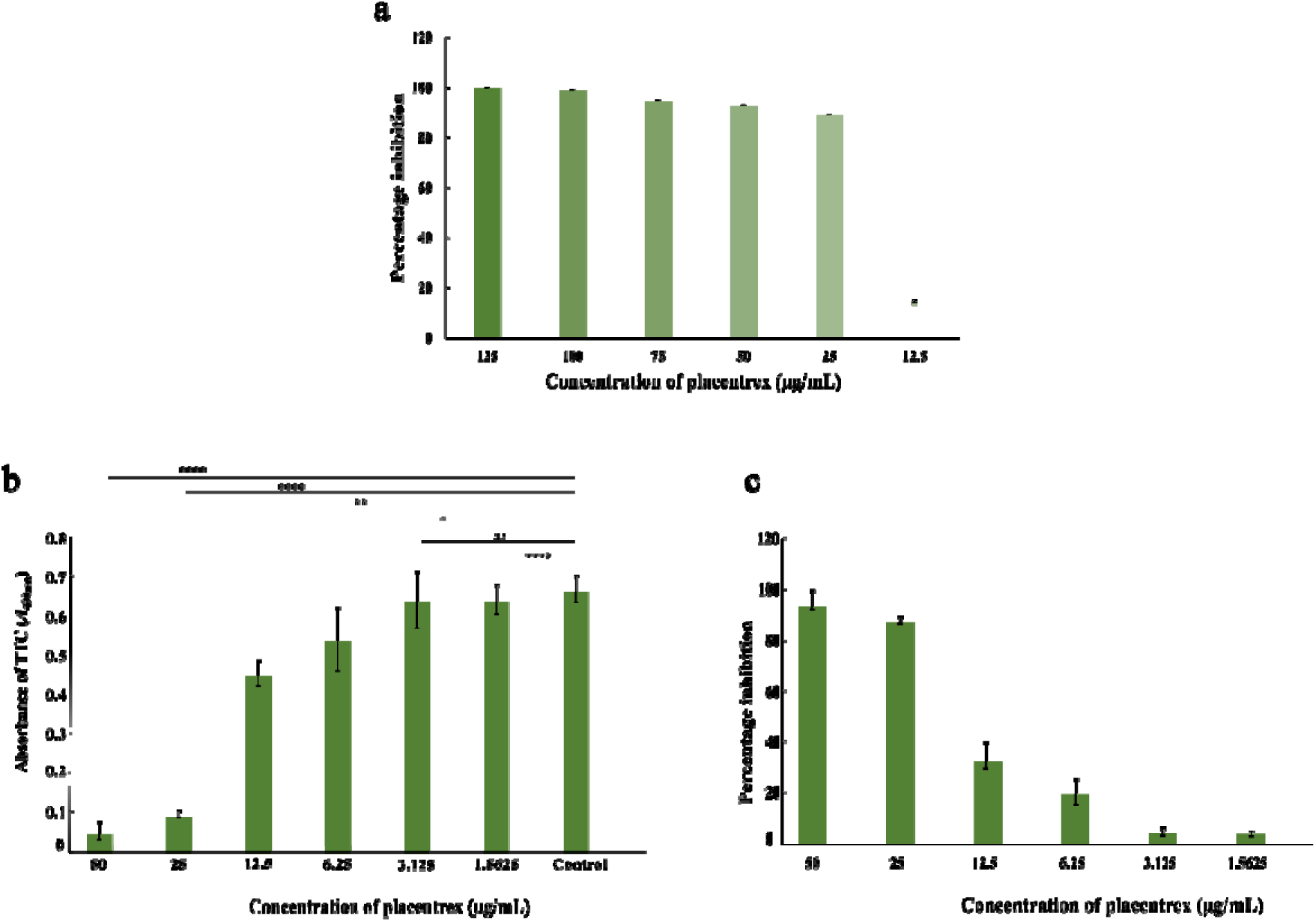
a) Percentage inhibition of preformed biofilm by PA cells in the presence of different concentrations of Placentrex by the Crystal Violet staining method, b) Quantitative assessment of the **v**iability of biofilm bacteria by the TTC assay, c) Percentage inhibition of viable cells within the biofilm in the presence of different concentrations of Placentrex by the TTC assay

This reduction in biofilm biomass prompted further investigation into whether Placentrex also affects the viability of bacterial cells embedded within the biofilm. Assessing cell viability is essential to determine if the observed biofilm clearance reflects true bacterial killing rather than mere structural disruption. Because the bacterial cells in the biofilms tend to re-grow in the presence of favourable conditions. Therefore, we analysed the effect of Placentrex on the viability of bacterial cells within the biofilm. Interestingly, the biofilm viability was reduced by approximately 93% and 87% upon treatment with 50 µg/mL and 25 µg/mL of Placentrex, respectively (Fig. 2b and 2c).

### Placentrex reduced the ratio of live/dead cells in PA biofilm

Here we observed that with increasing concentration of Placentrex, the proportion of dead cells (red fluorescence) increased significantly and the biofilms became thinner and looser compared to the control, demonstrating the accelerated death of bacterial cells within the biofilm. At the MIC concentration, both red and green fluorescence were extremely weak compared to the biofilm density in the control, indicating that most of the biofilm bacteria had been killed. Taken together, this demonstrates that Placentrex could change the permeability and integrity of the bacterial cell membrane, leading to the efficient eradication of the established PA biofilms (Fig. 3).

**Figure 3.**
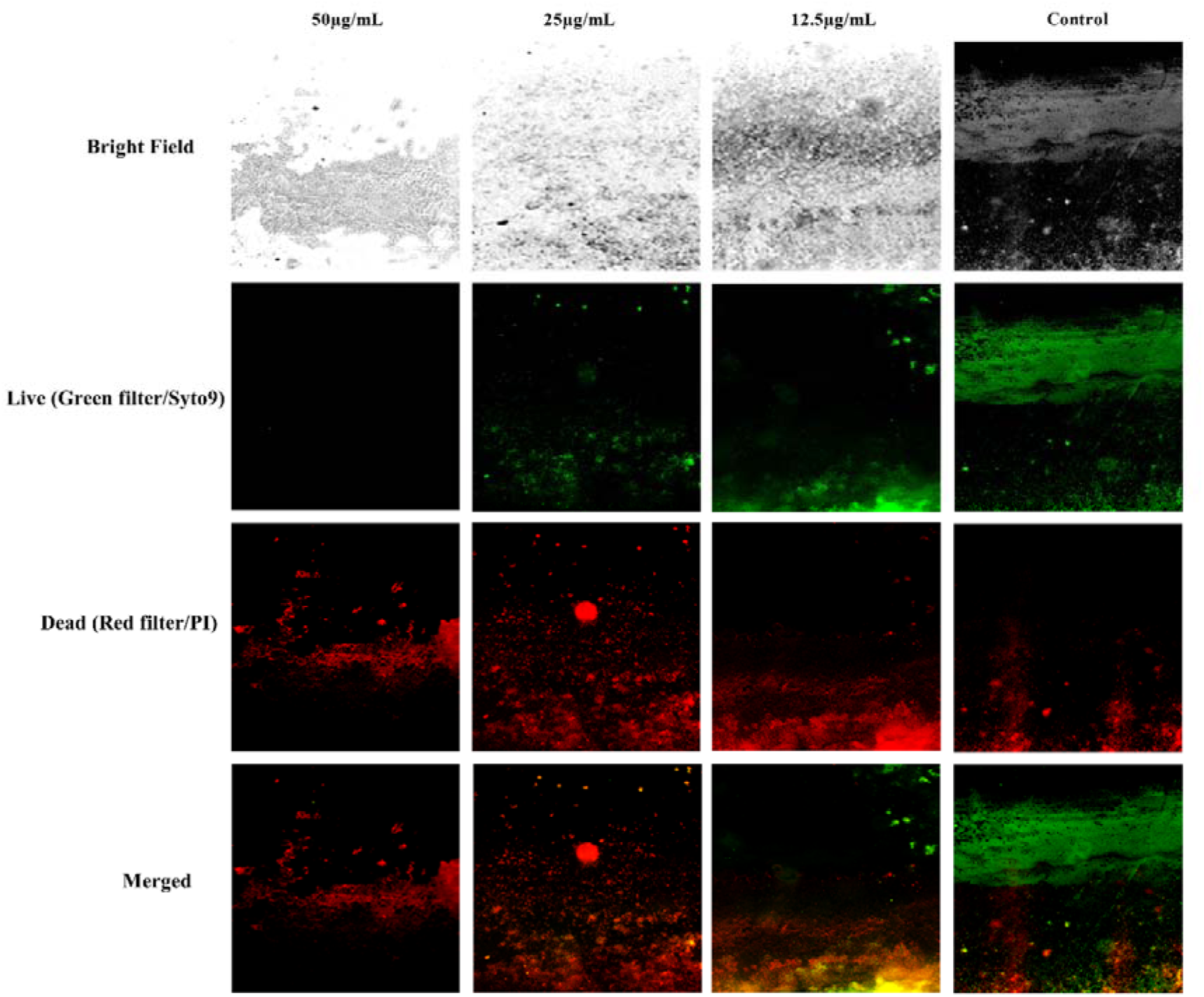
Fluorescence microscopy visualisation of biofilm formed by PA in the presence of different concentrations of placentrex

### Placentrex reduced the motility of PA

Bacterial motility is one of the key features that enable bacteria to move from one place to another in response to nutrients, chemical agents, or to transport themselves. While flagellar motility is a primary mechanism for movement that facilitates initial cell-surface contacts, which are essential for irreversible adhesion and the initiation of biofilm formation, other forms of motility, like swimming and twitching, can also contribute to biofilm formation. Hence, we studied the change in motility of PA upon Placentrex treatment. The zone of motility was visualised as a change in colour due to the reduction of TTC dye from the point of inoculation and represented as a change in the diameters across the zones of bacterial growth. Both twitching and swimming motility assays revealed significantly reduced motility for the PA as the placentrex concentration increases, following the same pattern as that in the biofilm inhibition assay. No movement from the site of inoculation was observed with 25µg/mL of Placentrex. The mean diameters of twitching motility (Fig S2a, S2b) were 35.7mm and 15.3mm, respectively, on control plates and 12.5 µg/mL of Placentrex-containing plate. The mean diameters of swimming motility (Fig. S2c, S2d) were 36.2 mm and 17.1 mm, respectively, on control plates and on plates containing 12.5 µg/mL Placentrex.

### Placentrex does not inhibit biofilm formation by reducing bacterial adhesion to the surface

The reversible attachment of bacterial cells to the surface is the first step of biofilm formation. Cell surface hydrophobicity (CSH) plays a significant role in influencing the adhesion properties of the bacteria (Vandana & Das, 2023). Hence, we analysed the adhesion of bacterial cells to the surface by MATH assay and observed that there is no considerable change in bacterial adhesion (Table S1). This demonstrates that inhibition of biofilm formation is not through the reduction in adhesion to the surface.

### Placentrex significantly reduced the EPS content of PA

The composition of the extracellular polymeric substance varies among bacterial species, particularly the polysaccharide content, which in turn determines the structural stability of the resultant biofilms. Therefore, we studied EPS from placentrex-treated samples. It was observed that the isolated EPS samples were insoluble in water or PBS (Phosphate-Buffered Saline) but readily dissolved in 0.1 M NaOH. This indicates that the matrix has strong intermolecular interactions with high protein or cross-linked polysaccharide content (Di Martino, 2018; Melo et al., 2022). Following isolation, the total protein and sugar content in 1mg/mL of all EPS samples were estimated using the Bradford assay and Phenol-Sulphuric Acid method, respectively (Table 1a). It was observed that the total sugar concentration decreased by approximately 17.29% to 54.9% as the Placentrex concentration increased. Similarly, the total protein in the EPS also reduced drastically from 18.5% to 92.6% with increasing concentration of Placentrex. These results also support the speculation that the matrix of this PA is rich in protein (Di Martino, 2018; Melo et al., 2022).

**Table 1a.**
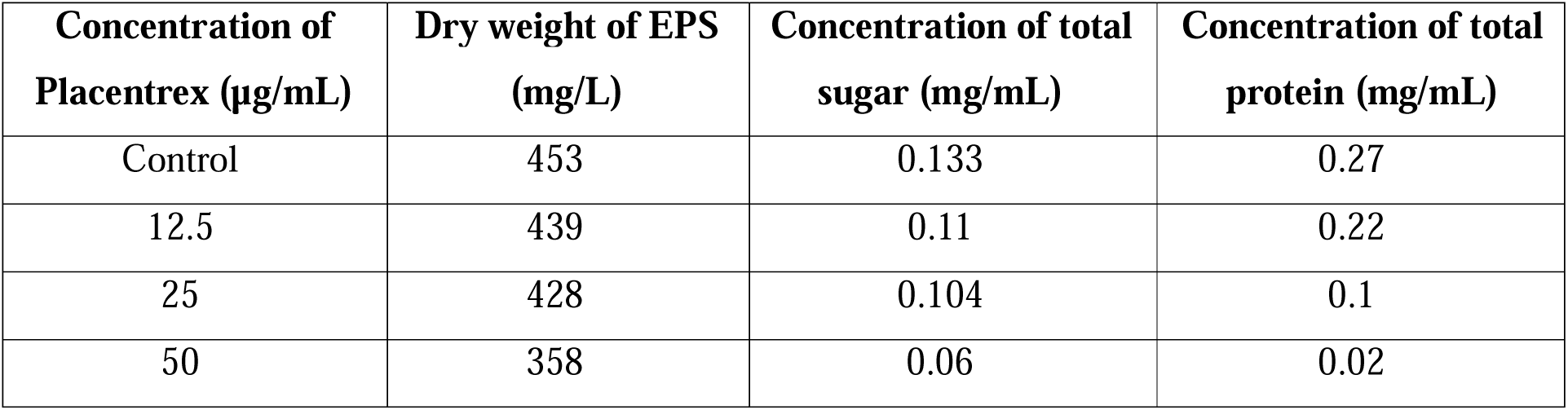
Change in concentration of total sugar and protein in PA EPS on placentrex treatment.

| Concentration of Placentrex ($\mu\text{g/mL}$) | Dry weight of EPS (mg/L) | Concentration of total sugar (mg/mL) | Concentration of total protein (mg/mL) |
| --- | --- | --- | --- |
| Control | 453 | 0.133 | 0.27 |
| 12.5 | 439 | 0.11 | 0.22 |
| 25 | 428 | 0.104 | 0.1 |
| 50 | 358 | 0.06 | 0.02 |

Further, the crude EPS was passed through Sepharose CL-6B gel filtration chromatography (GFC) column, and the total sugar concentration from each eluted tube was analysed by phenol-sulphuric acid assay to determine the sugar composition and molecular weight of EPS polysaccharides. Thin-layer chromatography (TLC) was performed to visualise the sugar composition of each eluted tube (Fig. S3a-d). The chromatogram derived from GFC revealed a single, well-defined peak (Fig. S4a- d). This suggests that the EPS of PA consists mainly of a single larger polysaccharide that constitutes the matrix along with other components, and the average molecular weight of this polysaccharide was estimated to be ∼1062 kDa using a standard dextran curve (Fig. S5).

Further, the eluted fractions containing the polysaccharide were pooled and analysed for the total polysaccharide concentration by phenol-sulphuric acid assay. This pooled fraction was then digested and neutralised, and subsequently visualised its monomeric sugars by TLC and analysed its composition by HPLC. Each EPS hydrolysate was visualised by running against different standard sugars on a TLC plate, showing four bands with retention factor (R_f_) values of 0.09, 0.21, 0.4, and 0.76, respectively. But all the samples showed one prominent band with an R_f_ value of 0.4, corresponding to the standard glucose R_f_ value (Fig. 4). To clarify the TLC data, the hydrolysed EPS samples were subsequently fractioned by HPLC against different monosaccharide standards. Consistent with the TLC profile, all HPLC chromatograms showed a single peak (Fig. S6). The concentration of monosaccharides in each exopolysaccharide was analysed by calculating the area under the peak compared with the peak area of the standard. It was observed that, across different Placentrex concentrations, the monosaccharide concentration was almost the same as the EPS-containing sugar concentration before hydrolysis (Table 1b). This suggests that the monosaccharide composition of EPS samples does not alter with different concentrations of Placentrex.

**Figure 4:**
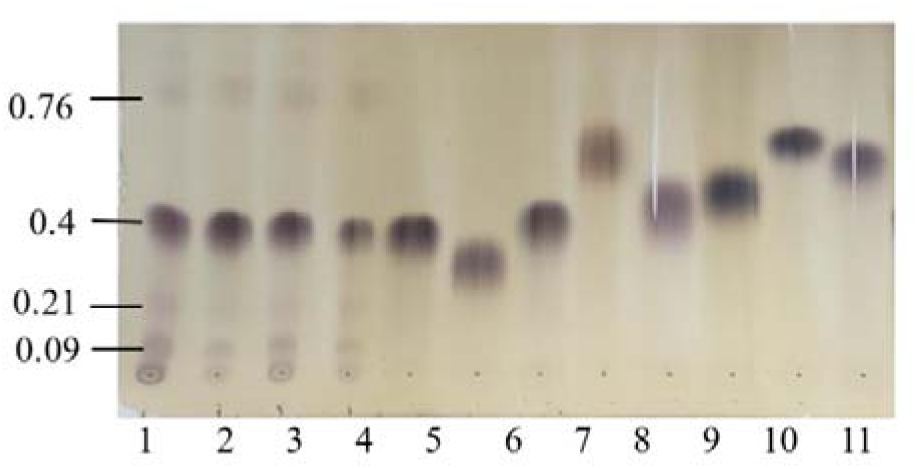
TLC profile of EPS hydrolysate. 1: control, 2: 12.5µg/mL placentrex treatment, 3: 25µg/mL Placentrex treatment, 4: 50µg/mL placentrex treatment) and different monosaccharide standards (5: glucose, 6: galactose, 7: fructose, 8: fucose, 9: mannose, 10: arabinose, 11: xylose, 12: ribose

**Table 1b.** Monosaccharide profiles of PA EPS by HPLC following hydrolysis.

| Concentration of Placentrex (µg/mL) | Concentration of EPS before hydrolysis (mg/ml) | Peak area of monosaccharide after hydrolysis of EPS in the HPLC chromatogram at ≈11.97 min [nRIU*S] | Concentration of total sugar (mg/mL) |
| --- | --- | --- | --- |
| Control | 5.2 | $1.97 \times 10^6$ | 5.02 |
| 12.5 | 4.5 | $1.572 \times 10^6$ | 4.01 |
| 25 | 4 | $1.537 \times 10^6$ | 3.92 |
| 50 | 2.2 | $8.587 \times 10^5$ | 2.19 |

### Placentrex alters the transcriptional landscape of PA to suppress biofilm

To elucidate the transcriptional response of PA to Placentrex, RNA sequencing was performed on bacterial cells treated with 12.5 µg/mL of Placentrex for 16 hours. Comparative transcriptomic analysis identified 134 differentially expressed genes (DEGs) (log FC ≥ 2) relative to the untreated control, of which 73 were up-regulated, and 61 were down-regulated (Fig 5; Tables S2 and S3). The greater number of up-regulated genes relative to down-regulated genes indicates a broad and active transcriptional response by PA upon exposure to Placentrex. Among the notably down-regulated genes were the GntP family permease, extracellular solute-binding protein, type 4b pilus Flp major pilin, urease accessory protein UreE, AraC family transcriptional regulator, aromatic amino acid permease, and an MFS transporter previously implicated in biofilm formation, collectively suggesting disruption of multiple cellular functions critical to biofilm establishment.

**Figure 5.**
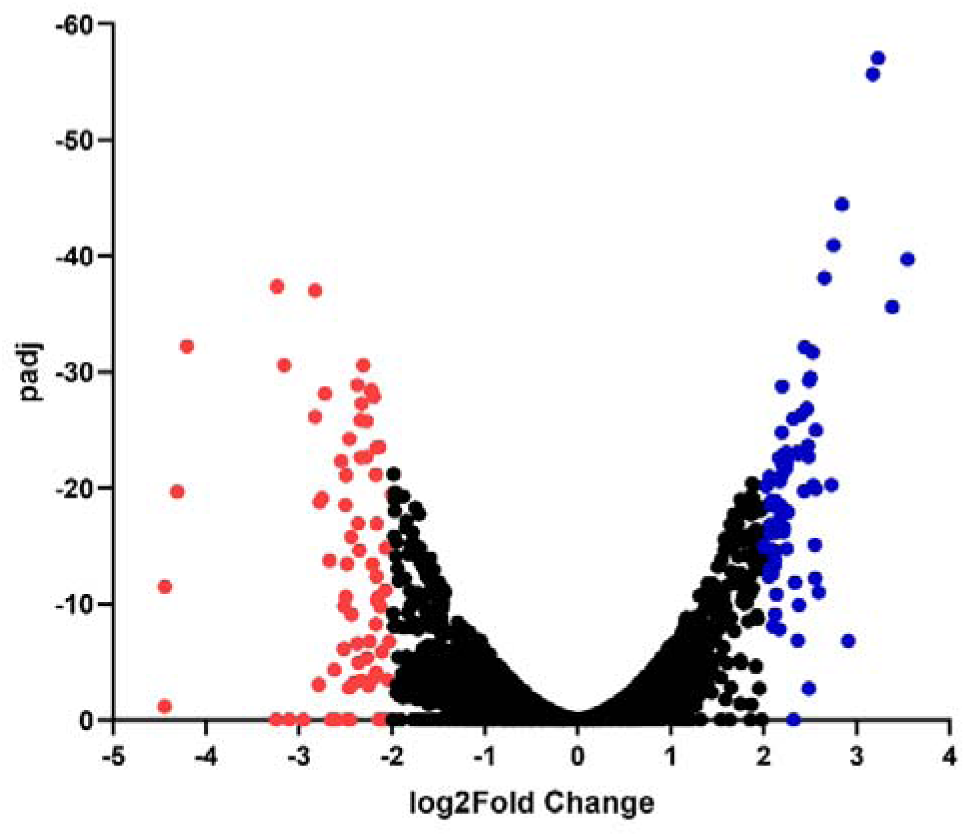
Volcano plot of differential gene expression between placentrex treatment and control groups. The result showed 73 genes up-regulated and 61 genes down-regulated in Placentrex- treated PA compared to untreated. Differentially expressed genes were selected at a fold change ≥ ±2 and FDR ≤ 0.05. The red colour indicates downregulated genes, while the blue colour indicates up-regulated genes.

Gene Ontology (GO) annotation of the DEGs (Table S4) across the three functional categories: biological process (BP), cellular component (CC), and molecular function (MF) further substantiated these findings (Fig. S7). Within the CC category, genes associated with the cytoplasm were predominantly down-regulated. In the BP category, significant down-regulation was observed in genes governing amino acid catabolic processes, including proteinogenic amino acid catabolism, L-amino acid catabolism, aromatic amino acid metabolism, aromatic amino acid family catabolic processes, and L-phenylalanine metabolism. In the MF category, genes involved in transferase and catalytic activity were similarly down-regulated. Additionally, within the CC category, genes associated with the plasma membrane, cell periphery membrane, intracellular anatomical structures, and cellular anatomical structures were found to be down-regulated, which are processes closely linked to the quorum sensing (QS) regulatory network and biofilm architecture in PA. Collectively, these findings suggest that Placentrex perturbs key metabolic pathways, thereby impeding the capacity of PA to form and maintain biofilms.

## Discussion

Biofilm formation is one of the principal mechanisms of virulence in Gram- positive and Gram- negative pathogens. Biofilm-associated infections are chronic and recurrent, as biofilm promotes bacterial survival within the host. *P. aeruginosa* is one of the major pathogens involved in opportunistic infections in humans, and biofilm formation complicates this further by facilitating bacterial colonisation, enhancing antimicrobial resistance, and evading the host immune system (Elmanama et al., 2020; Maurice et al., 2018; Tuon et al., 2022). Consequently, traditional antibiotics demonstrated limited effectiveness in preventing biofilm formation and treating biofilm-associated infections, making the development of innovative antimicrobial agents and biofilm-disrupting therapeutic approaches essential for controlling *P. aeruginosa* infections. In a preliminary study, human placental extract (HPE) containing fibronectin type III (FN3)-like peptide and Ubiquitin-like peptide has shown promising antibiofilm activity. HPE has been shown to inhibit quorum-sensing-dependent biofilm formation in both Gram-positive and Gram-negative organisms(Goswami et al., 2017).

The present study reveals that Placentrex is highly effective in combating biofilm and biofilm bacteria. To begin with, we studied the susceptibility of PA to Placentrex. We found Placentrex demonstrated substantial antibacterial and antibiofilm activity against PA, achieving bactericidal effects with complete eradication within 24 hours at 50µg/mL. This rapid action is consistent with a multifaceted mechanism of action, and to delineate the molecular basis of this activity, transcriptomic profiling (RNA-seq) was performed on Placentrex-treated PA. The RNA-seq analysis revealed a coordinated downregulation of genes involved in nutrient transport, pilus assembly, metabolic regulation, virulence enzyme maturation, and membrane efflux, which mechanistically explains the broad antibiofilm and antibacterial effect. Such polypharmacological transcriptional reprogramming reduces selective pressure for resistance compared to the single-target antibiotics and is consistent with the low resistance development potential (Baquero & Levin, 2021). Consistent with the above finding of bacterial killing, Placentrex significantly reduced the biofilm-forming ability of PA at MIC, MIC/2, and MIC/4, respectively, even after 72 hours of incubation. The two major steps in *P. aeruginosa* biofilm development are cell-surface attachment and cell-cell interactions(Qu et al., 2016). The RNA-seq data provide a precise molecular explanation for the disruption of the very first of these steps: the downregulation of the flp major pilin gene, which encodes the structural subunit of the Type IVb Flp pilus. Flp pili are required for adhesion to abiotic surfaces and eukaryotic cells. This process of attachment represents a critical mediator of irreversible surface attachment that transits planktonic bacteria into sessile biofilm communities (De Bentzmann et al., 2006). Downregulation of flp by Placentrex is therefore predicted to impair the initial colonisation step, restricting biofilm establishment at its earliest stage.

Notably, the MATH (Microbial Adhesion to Hydrocarbons) assay demonstrated no measurable effect on bacterial cell- surface hydrophobicity following Placentrex treatment. This finding is not contradictory to the observed anti-adhesion activity, but is in fact mechanistically informative. Flp/type IVb pili mediate adhesion through specific receptor-ligand interactions. Their hydrophobic N-terminal domain is sequestered within the pilus shaft core and does not contribute to bulk surface hydrophobicity (Alteri et al., 2022; Craig et al., 2004). The MATH assay measures non-specific hydrophobic partitioning driven principally by outer membrane lipid composition and LPS character, neither of which was significantly altered by Placentrex (Rosenberg, 2006; Z. Wang et al., 2015). The anti-adhesion effect of Placentrex is therefore speculated to be mediated by targeted transcriptional silencing of the Flp pilin assembly apparatus, and not by broad membrane-level hydrophobicity changes.

Preformed biofilms are mature, established communities of microorganisms embedded in a self-produced matrix on biotic or abiotic surfaces, exhibiting up to 1000-fold higher resistance to antimicrobial agents compared to planktonic cells. These established structures represent a major challenge in chronic, persistent infections and industrial contamination due to their recalcitrance to eradication(Oluwole, 2022). Preformed biofilms can be effectively dismantled through the targeted disruption of cell-cell adhesion, causing bacterial aggregates to separate and bacteria to revert to their free-floating planktonic form(Melia et al., 2021). Placentrex eradicated the preformed PA biofilm by approximately 93% and 89% at 50 µg/mL and 25 µg/mL, respectively, after 24 hours. The RNA-seq transcriptomic data reveal a mechanistic basis for this striking eradication efficiency, beginning with the downregulation of an extracellular solute-binding protein (SBP). Extracellular solute binding protein or SBP is the periplasmic ligand-sensing subunit of an ABC-type transport system. In PA, the substrate-binding protein DppA1 of the DppBCDF ABC transporter has been directly demonstrated to be essential for biofilm formation and structural maintenance and its inactivation results in 68-fold less biofilm in static models and complete abolition of biofilm in flow cells (Lee et al., 2018). In addition, it can sense the external nutrient availability, linking the SBP to the maintenance of preformed biofilm (Lee et al., 2018). Placentrex-mediated downregulation of the extracellular solute binding protein therefore mimics a nutrient-depleted signalling state, deactivating the bacterium’s nutrient-sensing ability, thus compromising the biofilm maintenance programme and eventually destabilising the preformed biofilm from within.

Adding to this nutrient-sensing disruption, RNA-seq identified downregulation of an AraC family transcriptional regulator, a master regulator of quorum sensing (QS) and virulence gene expression in PA. For example, a major transcriptional regulator belonging to the AraC-type family, VqsM, acts as a global master regulator that controls the production of quorum-sensing (QS) signalling molecules (N-acylhomoserine lactones) and extracellular virulence factors in PA(Dong et al., 2005). QS-driven gene expression is indispensable for the maintenance and structural integrity of mature biofilms. Downregulation of the AraC family transcriptional regulator by Placentrex can therefore be predicted to collapse the QS network that sustains preformed biofilm cohesion, simultaneously reducing matrix structuring and the expression of genes required to maintain cell-cell connections within the established biofilm.

The simultaneous downregulation of an MFS (Major Facilitator Superfamily) transporter further compounds the disruption of preformed biofilm maintenance. In PA, active efflux is required for the export of the major QS autoinducer N-(3-Oxododecanoyl)-L-homoserine lactone, which cannot exit the cell efficiently by passive diffusion alone. Efflux-deficient mutants show strong intracellular accumulation of AHLs and markedly reduced biofilm formation (Alav et al., 2018; Pearson et al., 1999). Their downregulation impairs autoinducer export, effectively internalising QS signals and disrupting the extracellular concentration thresholds required for coordinated biofilm gene expression (Minagawa et al., 2012). Concurrently, reduced MFS efflux capacity lowers the bacterium’s intrinsic tolerance to antimicrobials and host defence compounds, rendering cells embedded within the preformed biofilm matrix more susceptible to Placentrex (Alcalde-Rico et al., 2020; Ciofu & Tolker-Nielsen, 2019). The multi-effect of impaired intercellular communication, nutrient unavailability to the biofilm core, and reduced efflux-mediated protection can therefore be speculated to create a permissive environment for biofilm eradication (Alcalde-Rico et al., 2020; Ciofu & Tolker-Nielsen, 2019).

At the matrix level, Placentrex induced significant structural modifications to the EPS of *P. aeruginosa* preformed biofilms. The increased solubility of treated EPS in 0.1M NaOH indicates enhanced cross-linking between matrix components mediated by carboxyl, phosphoric, or sulfhydryl groups (Melo et al., 2022), producing a matrix that is structurally rigidified but physiologically less adaptable analogous to the action of glycoside hydrolases like PelA and PslG, which strengthen matrix adhesion while simultaneously impairing the biofilm’s capacity to respond to environmental changes (Baker et al., 2016; Colvin et al., 2011). Gel filtration chromatography confirmed that Placentrex preserved the core polysaccharide molecular weight while altering its physiological behaviour, indicating disruption of non-covalent interactions rather than polymer fragmentation, a mechanism similar to that of curcumin, which alters EPS solubility without depolymerisation to enhance antibiotic penetration (Sharifian et al., 2020). The RNA-seq data provide a multi-layered transcriptional basis for the observed EPS structural changes, operating through three convergent mechanisms. First, downregulation of gntP reduces carbon flux through the Entner-Doudoroff pathway, limiting metabolic precursors for rhamnolipid biosynthesis. Rhamnolipids are critical biosurfactants that govern water channel formation and EPS matrix architecture in PA biofilms (Davey et al., 2003). Second, downregulation of the aromatic amino acid permease restricts the import of phenylalanine and tyrosine, thereby depleting the biosynthesis of exopolysaccharide, pyoverdine and reducing the metabolic capacity of biofilm cells to sustain matrix production (Palmer et al., 2010). Third, and directly relevant to EPS matrix assembly, MFS-type efflux pumps have been demonstrated to facilitate the export of biofilm matrix sugars, including glucose and arabinose, that are essential structural components of the EPS scaffold (Alav et al., 2018). The putative MFS transporter PA2114 of P. aeruginosa is up-regulated greater than 2.5- fold in developing biofilms compared with planktonic cells (Alav et al., 2018), suggesting that MFS-mediated sugar export is an active component of EPS matrix assembly. Downregulation of this MFS transporter by Placentrex is therefore predicted to impair the supply of matrix sugar components required for EPS structural maintenance, compounding the metabolic deficits arising from GntP and aromatic amino acid permease suppression. Together, these transcriptional changes produce an EPS that is structurally compromised and metabolically unsustainable.

The destabilisation of biofilm architecture was further confirmed by increased cell death following Placentrex treatment, as evidenced by TTC assay and Live/Dead staining. This cell death is attributable to the convergent effects of: (i) structural rigidification of the EPS matrix, which reduces nutrient and oxygen diffusion to deeper biofilm layers; (ii) collapse of the QS communication network via AraC regulator and MFS transporter downregulation, which disrupts coordinated metabolic adaptation within the biofilm; (iii) impaired nutrient sensing via SBP downregulation, which mimics starvation signalling and triggers dispersal-associated cell death in non-dispersing cells; and (iv) reduced carbon and amino acid import via GntP and aromatic amino acid permease downregulation, which starves metabolically active biofilm cells of essential substrates. The cumulative effect of matrix rigidification, together with metabolic substrate limitation and impaired EPS matrix supply, provides a mechanistically coherent explanation for the near-complete biofilm cell death observed experimentally.

## Conclusion

In conclusion, Placentrex is highly effective in combating PA biofilms and planktonic bacteria through a coordinated, multi-target mechanism of action that spans transcriptional, metabolic, structural, and virulence-associated pathways. The absence of MATH-detectable changes in cell surface hydrophobicity, far from being contradictory, confirms that these mechanisms operate through highly specific molecular interactions rather than broad membrane disruption. The integration of RNA-seq transcriptomic data with functional antibiofilm, EPS characterisation, MATH, TTC, and Live/Dead assay results provides a comprehensive picture. Placentrex simultaneously silences pilus-mediated attachment (Flp pilin), disrupts the carbon metabolic backbone fuelling biofilm architecture (GntP, aromatic AA permease), decouples environmental sensing from biofilm induction (solute-binding protein), dismantles QS-driven biofilm maintenance (AraC regulator, MFS transporter), and inactivates a urease virulence pathway with no human ortholog (UreE). These findings collectively support the use of Placentrex as a promising therapeutic agent for the management of biofilm-associated PA infections.

## Acknowledgement

We thank M/s Albert David Limited, Kolkata, India, for providing us with Placentrex.

## Author contributions

Bayomi Biju (Data curation, Formal analysis, Investigation, Methodology, Validation, Visualisation, Writing – original draft, Writing – review & editing), Tejavath Ajith ( Data curation, Formal analysis, Investigation, Methodology, Validation, Visualisation, Writing – original draft, Writing – review & editing), Ajit R Sawant (Data curation, Formal analysis, Investigation, Methodology, Validation, Visualisation, Writing – review & editing), Sachin Maji (Data curation, Formal analysis, Investigation, Methodology, Validation, Writing – review & editing), Tridib Neogi (Conceptualization, Funding, Investigation, Resources, Validation, Writing – review & editing), Piyali Datta Chakraborty (Conceptualization, Funding, Investigation, Resources, Validation, Writing – review & editing), Anindya Sundar Ghosh (Conceptualization, Funding acquisition, Investigation, Project administration, Resources, Supervision, Validation, Writing – original draft, Writing – review & editing)

## Data availability

All data generated or analysed during this study are included in this article and its supplementary section.

