## Supplementary File for "Placentrex disrupts the biofilm formation of *Pseudomonas aeruginosa* through multi-target transcriptional reprogramming"

**Title:**

Affiliations:

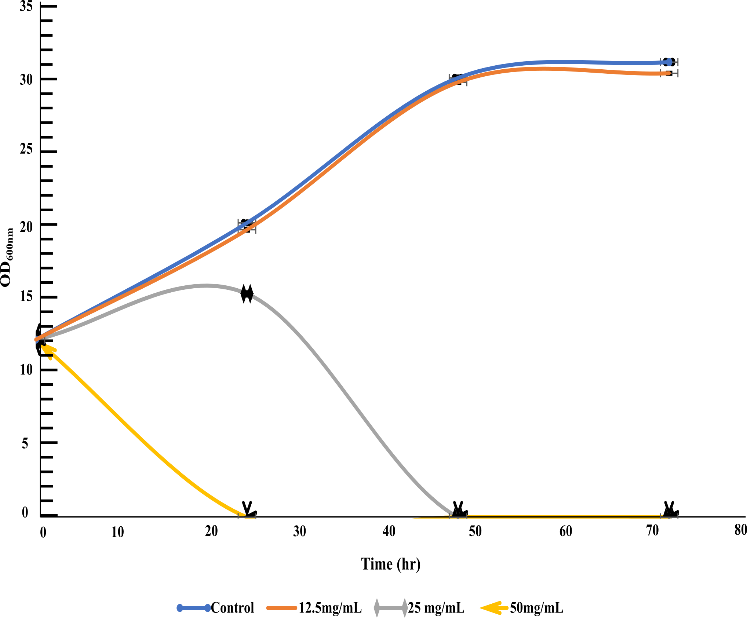

**Figure S1.** Time kill curve of PA treated with ¼ × MIC, ½ × MIC, and 1 × MIC of Placentrex.

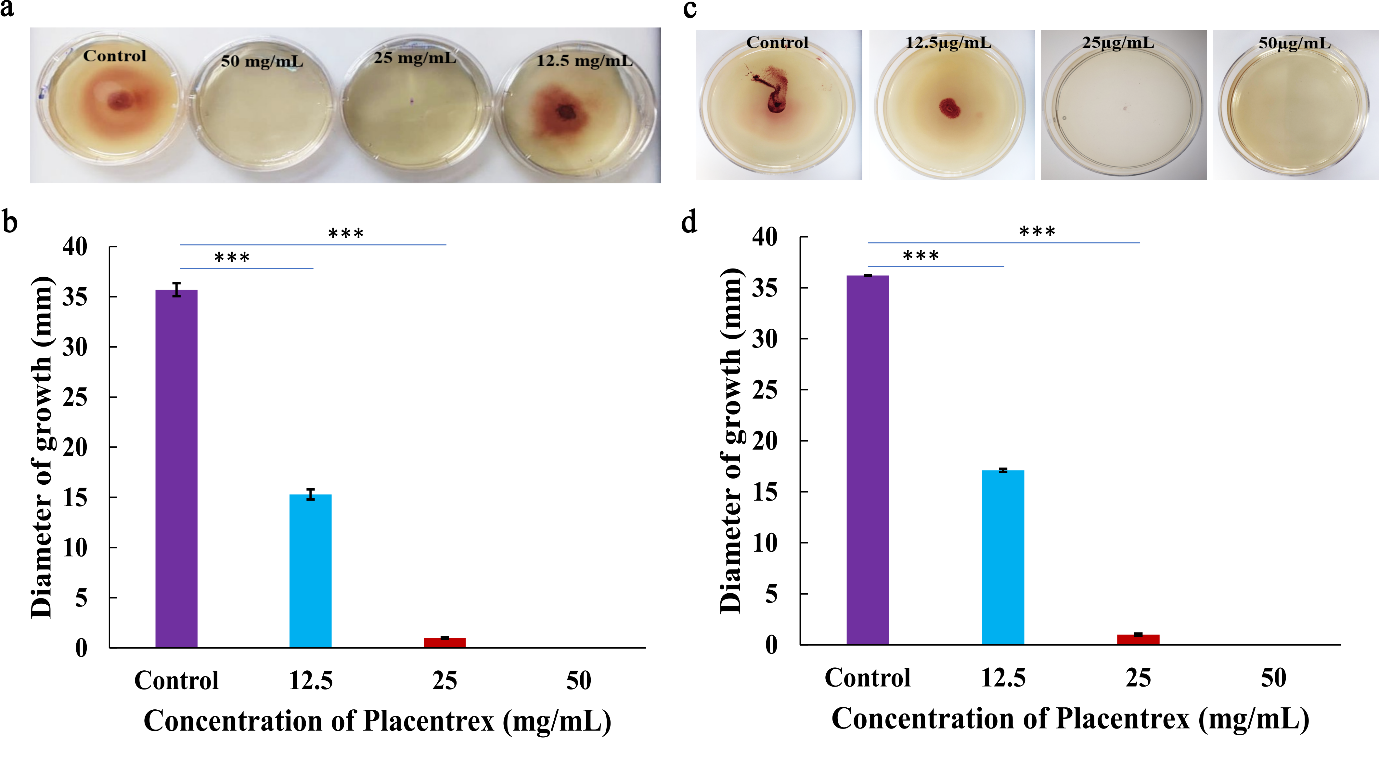

**Figure S2.** Motility assay for PA. a)Bacterial movements on twitching media, b)Measurement of the twitching motility, c)Bacterial movements on swimming media, d)Measurement of the swimming motility

**Table S1:** Change in percentage Hydrophobicity of PA on placentrex treatment

| **Placentrex Concentration (mg/mL)** | **%Hydrophobicity** |
| --- | --- |
| Control | 43.35 |
| 12.5 | 41.57 |
| 25 | 40.66 |

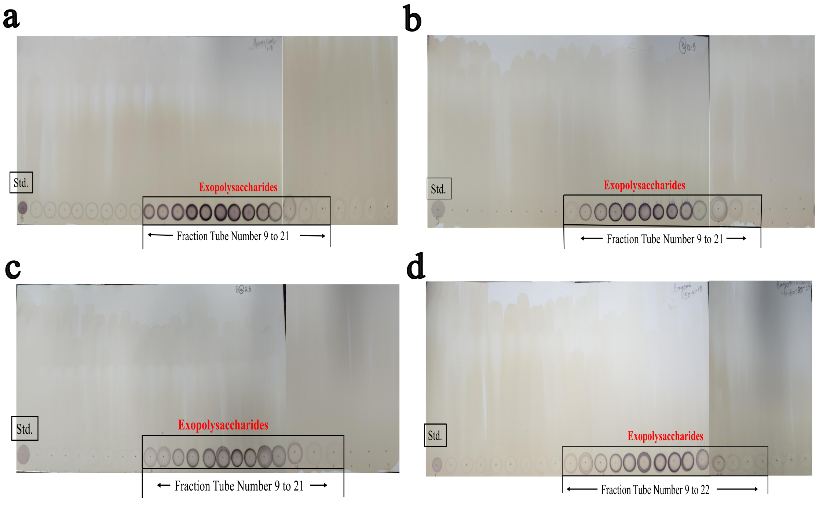

**Figure S3**: TLC profile of GFC eluted EPS. a: control, b: 12.5mg/mL placentrex treatment, c: 25mg/mL Placentrex treatment, d: 50mg/mL placentrex treatment. Std refers to Lipopolysaccharides from *Salmonella enterica* serotype enteritidis.

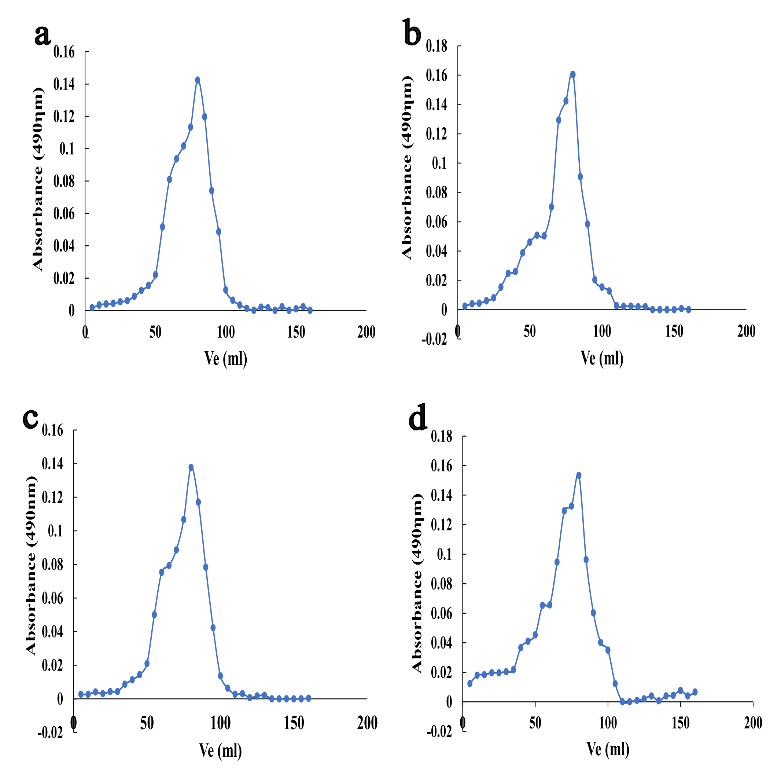

**Figure S4**: Gel filtration chromatography profile of EPS treated with different concentrations of Placentrex with respect to the untreated control. a: control, b: 12.5mg/mL placentrex treatment, c: 25mg/mL placentrex treatment, d: 50mg/mL placentrex treatment, Ve is the elution volume

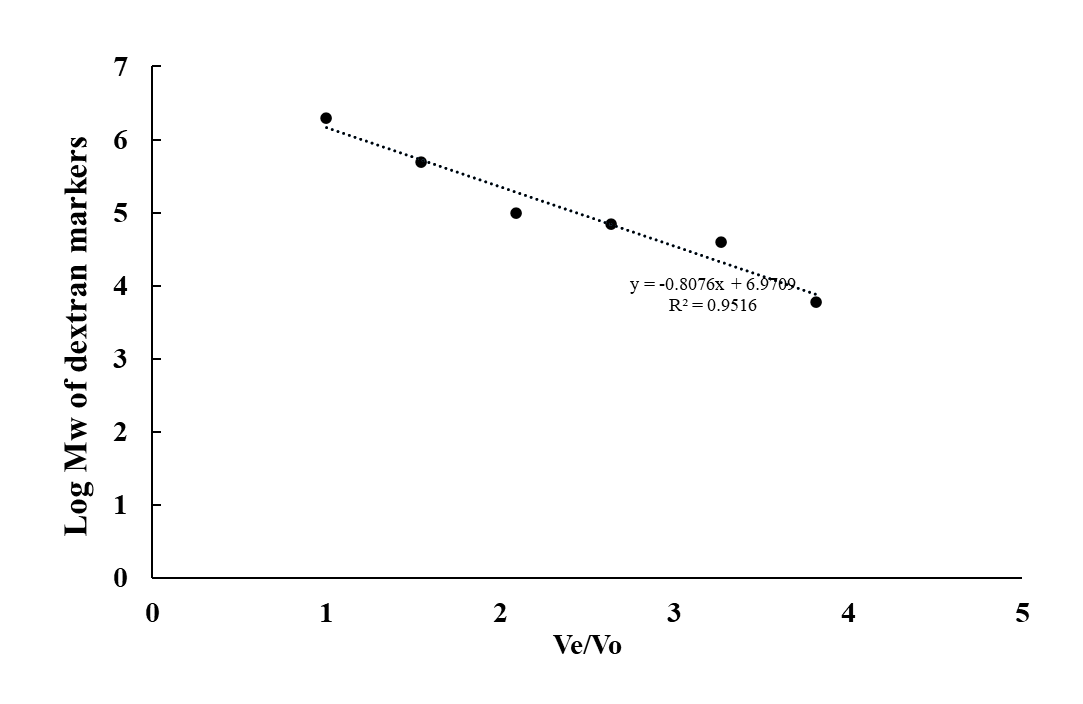

**Figure S5:** Standard curve of log Mw of dextran markers plotted against the ratio of eluted volume to void volume.

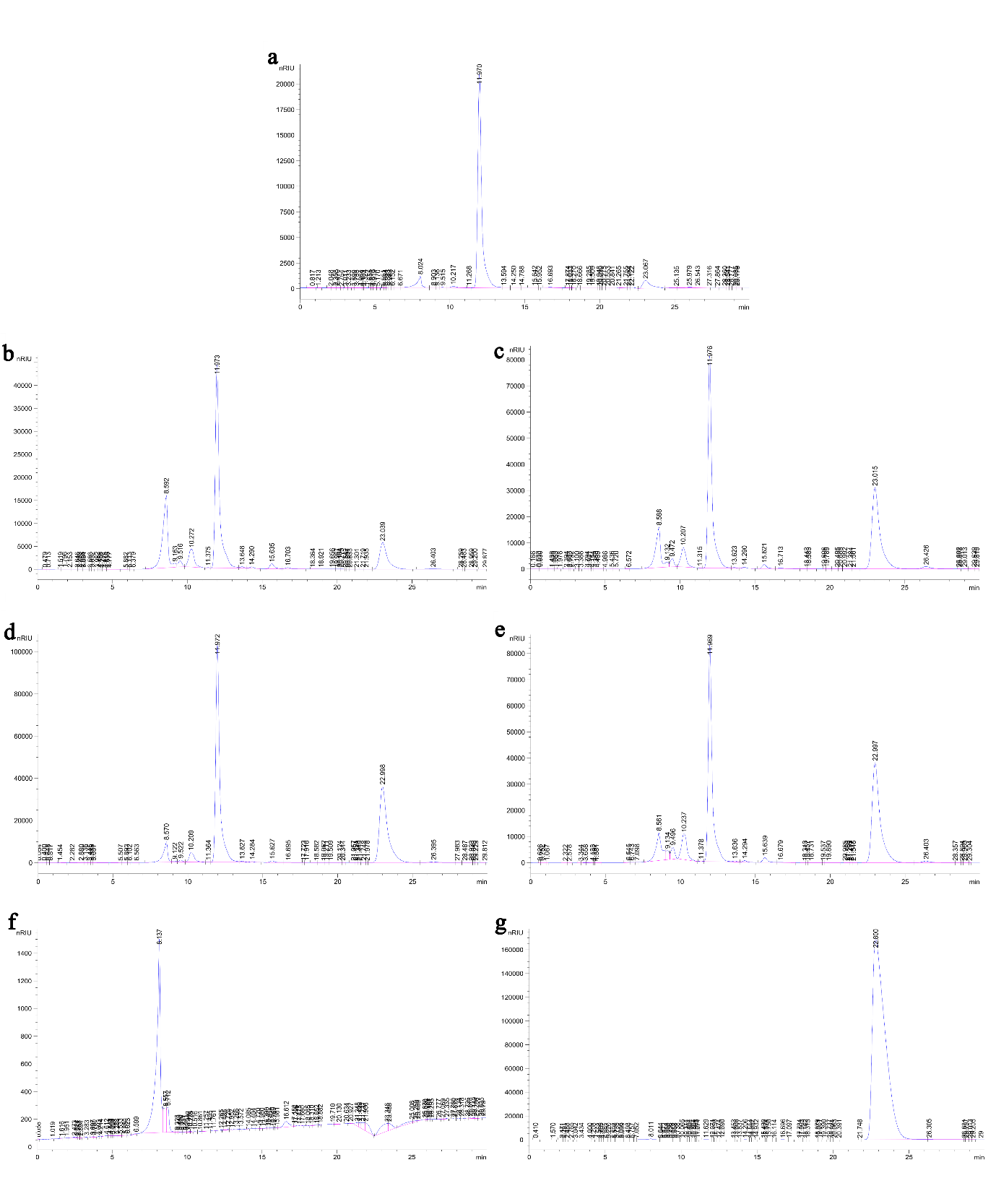

**Figure S6:** Monosaccharide composition analysis of EPS by HPLC. a) standard glucose, b) 50mg/mL treated EPS, c) 25mg/mL treated EPS, d) 12.5mg/mL treated EPS, e) Control, f) water, g) methanol.

**Table S2:** List of upregulated genes

| **Gene** | **log2FoldChange** | **Product** |
| --- | --- | --- |
|  | 3.548147201 | DUF1652 domain-containing protein |
|  | 3.382978121 | DUF1652 domain-containing protein |
|  | 3.229575835 | hypothetical protein |
|  | 3.177317772 | hypothetical protein |
|  | 2.91183154 | Coenzyme PQQ synthesis protein B |
|  | 2.842382553 | DUF485 domain-containing protein |
|  | 2.752890844 | Methyltransferase type 11 domain-containing protein |
|  | 2.727747883 | Putative major facilitator superfamily transporter |
|  | 2.65495853 | Methyltransferase type 11 domain-containing protein |
|  | 2.592307981 | Secreted protein |
|  | 2.560316704 | DUF1682 domain-containing protein |
| acs | 2.560062371 | Sel1 domain-containing protein |
|  | 2.553282834 | Putative major facilitator superfamily transporter |
|  | 2.551939142 | pyrroloquinoline quinone biosynthesis protein PqqB |
| nuoM | 2.533665172 | DUF485 domain-containing protein |
|  | 2.528766638 | Lipoprotein |
|  | 2.503081642 | L-lactate permease |
|  | 2.490488688 | Lipoprotein |
| lctP | 2.485744349 | TonB-dependent siderophore receptor |
| nuoN | 2.481667868 | LTXXQ domain protein |
|  | 2.474850089 | Sel1 domain-containing protein |
|  | 2.461687852 | hybrid sensor histidine kinase/response regulator |
| lldD | 2.439890493 | histidine kinase |
|  | 2.432539966 | hypothetical protein |
| nuoH | 2.402729204 | hypothetical protein |
|  | 2.378207606 | TonB-dependent siderophore receptor |
| nuoH | 2.372601118 | UPF0337 protein PA4738 |
| nuoL | 2.365949532 | Secreted protein |
| lldD | 2.362741688 | UPF0337 protein PA4738 |
| nuoJ | 2.36040362 | lactate permease LctP family transporter |
| carO | 2.3355754 | pyrroloquinoline quinone biosynthesis peptide chaperone PqqD |
|  | 2.318420162 | NADH-quinone oxidoreductase subunit M |
| carO | 2.259547688 | putative membrane transporter protein |
|  | 2.253052366 | hypothetical protein |
|  | 2.246736074 | pyrroloquinoline-quinone synthase PqqC |
| actP | 2.246361379 | NADH-quinone oxidoreductase subunit NuoH |
|  | 2.242174845 | NADH-quinone oxidoreductase subunit NuoN |
|  | 2.240321274 | FMN-dependent L-lactate dehydrogenase LldD |
|  | 2.216648572 | Yip1 domain-containing protein |
|  | 2.209275588 | NADH dehydrogenase I chain L |
|  | 2.200482327 | NADH-quinone oxidoreductase subunit L |
| piuC | 2.197971073 | FMN-dependent L-lactate dehydrogenase LldD |
| nuoI | 2.197816079 | acetate--CoA ligase |
| nuoI | 2.193172449 | Methyl-accepting chemotaxis protein |
|  | 2.190054075 | NADH-quinone oxidoreductase subunit J |
| pqqE | 2.184664564 | NADH-quinone oxidoreductase subunit NuoI |
| pigA | 2.178576591 | pyrroloquinoline quinone biosynthesis protein PqqE |
|  | 2.173045423 | BON domain-containing protein |
|  | 2.167141802 | hypothetical protein |
|  | 2.165652873 | NADH-quinone oxidoreductase subunit NuoH |
|  | 2.137761492 | Inner membrane protein YohK |
| pqqB | 2.125081695 | NADH-quinone oxidoreductase subunit NuoK |
|  | 2.123950806 | (Fe-S)-binding protein |
|  | 2.123757069 | BON domain-containing protein |
| pqqC | 2.12319044 | Cytochrome b561 bacterial/Ni-hydrogenase domain-containing protein |
|  | 2.123142408 | Iron-uptake factor PiuC |
| nuoK | 2.116557563 | NADH-quinone oxidoreductase subunit NuoI |
|  | 2.115156543 | UPF0391 membrane protein PLES_58781 |
|  | 2.104293705 | NADH-quinone oxidoreductase |
|  | 2.09936678 | acetyl-CoA hydrolase/transferase C-terminal domain-containing protein |
| nuoK | 2.093720265 | NADH-quinone oxidoreductase subunit NuoK |
|  | 2.088457468 | Succinyl-CoA:coenzyme A transferase |
|  | 2.085517012 | Cytochrome b561 bacterial/Ni-hydrogenase domain-containing protein |
| pqqD | 2.078381772 | Fe2+-dependent dioxygenase |
|  | 2.070528651 | biliverdin-producing heme oxygenase PigA |
| yohK | 2.070330444 | Yip1 domain-containing protein |
|  | 2.069535753 | (Fe-S)-binding protein |
| piuC | 2.062725217 | Calcium-regulated OB-fold protein CarO |
|  | 2.056152426 | UPF0391 membrane protein PLES_58781 |
|  | 2.039283809 | Calcium-regulated OB-fold protein CarO |
|  | 2.024957831 | Cation/acetate symporter ActP |
|  | 2.021106908 | putative membrane transporter protein |
|  | 2.004681725 | aminotransferase class V-fold PLP-dependent enzyme |

**Table S3:** List of downregulated genes

| **Gene** | **log2FoldChange** | **Product** |
| --- | --- | --- |
|  | -2.000589994 | glycine cleavage system aminomethyltransferase GcvT |
| cioA | -2.029586981 | AraC family transcriptional regulator |
| gntP | -2.045798938 | Amino acid permease/ SLC12A domain-containing protein |
| mmsA | -2.064845603 | DUF1329 domain-containing protein |
|  | -2.06589559 | Re/Si-specific NAD(P)(+) transhydrogenase subunit alpha |
| mmsB | -2.105497456 | Aromatic amino acid permease |
| phhC | -2.123626412 | type 4b pilus Flp major pilin |
|  | -2.135008876 | phenylalanine 4-monooxygenase |
|  | -2.147205715 | Secreted protein |
| phhC | -2.159602056 | type 4b pilus Flp major pilin |
| mmsB | -2.160857029 | 3-hydroxybutyrate dehydrogenase |
|  | -2.164976599 | phenylalanine 4-monooxygenase |
|  | -2.166030672 | Re/Si-specific NAD(P)(+) transhydrogenase subunit alpha |
|  | -2.169061195 | MFS transporter |
|  | -2.172025988 | Uncharacterized protein PA3568 |
| phhA | -2.172422877 | putative hydro-lyase |
| phhA | -2.188958078 | Aromatic-amino-acid aminotransferase |
|  | -2.18977517 | hypothetical protein |
|  | -2.205771528 | Amino acid permease/ SLC12A domain-containing protein |
|  | -2.219637142 | Aromatic-amino-acid aminotransferase |
|  | -2.222591619 | aldehyde dehydrogenase family protein |
|  | -2.237655266 | urease accessory protein UreE |
| gntP | -2.24794783 | hypothetical protein |
| gcvT | -2.265568802 | CoA transferase |
|  | -2.270904847 | Pterin-4-alpha-carbinolamine dehydratase |
|  | -2.272373289 | Uncharacterized protein PA3568 |
|  | -2.30655967 | Methylmalonate-semialdehyde dehydrogenase [acylating] |
|  | -2.323132123 | 3-hydroxyisobutyrate dehydrogenase |
| bdhA | -2.330865483 | urease accessory protein UreF |
|  | -2.332164588 | DUF3509 domain-containing protein |
|  | -2.337266902 | Pterin-4-alpha-carbinolamine dehydratase |
|  | -2.348858731 | Extracellular solute-binding protein |
| mdcA | -2.359838121 | hypothetical protein |
|  | -2.362354819 | DUF2845 domain-containing protein |
|  | -2.369459924 | 3-hydroxyisobutyrate dehydrogenase |
|  | -2.372567783 | 3-oxoacyl-[acyl-carrier-protein] reductase FabG |
|  | -2.400087754 | hypothetical protein |
|  | -2.434126047 | DUF1329 domain-containing protein |
| mdcA | -2.435679622 | Extracellular solute-binding protein |
|  | -2.456229189 | DUF3509 domain-containing protein |
| flp | -2.459281258 | DUF2790 domain-containing protein |
|  | -2.483401769 | Secreted protein |
| flp | -2.492358746 | DUF1654 domain-containing protein |
|  | -2.496345084 | malonate decarboxylase subunit alpha |
|  | -2.498725671 | ABM domain-containing protein |
| ureE | -2.512516987 | putative hydro-lyase |
| araC | -2.513382868 | L-serine ammonia-lyase, iron-sulfur-dependent, subunit alpha |
| fabG | -2.544635477 | DUF1654 domain-containing protein |
|  | -2.618184249 | ubiquinol oxidase subunit II |
|  | -2.668972877 | malonate decarboxylase subunit alpha |
|  | -2.719612173 | (S)-ureidoglycine aminohydrolase cupin domain-containing protein |
|  | -2.750221508 | Cyanide insensitive terminal oxidase, subunit III |
| cyoA | -2.775371741 | Cyanide insensitive terminal oxidase, subunit III |
|  | -2.787148165 | hypothetical protein |
|  | -2.823928315 | cyanide-insensitive cytochrome bd quinol oxidase subunit I |
|  | -2.824516458 | (S)-ureidoglycine aminohydrolase cupin domain-containing protein |
| ureF | -3.158442944 | FAD dependent oxidoreductase domain-containing protein |
|  | -3.232069946 | FAD dependent oxidoreductase domain-containing protein |
|  | -4.206388693 | GntP family permease |
|  | -4.307592151 | GntP family permease |
|  | -4.441047756 | hypothetical protein |

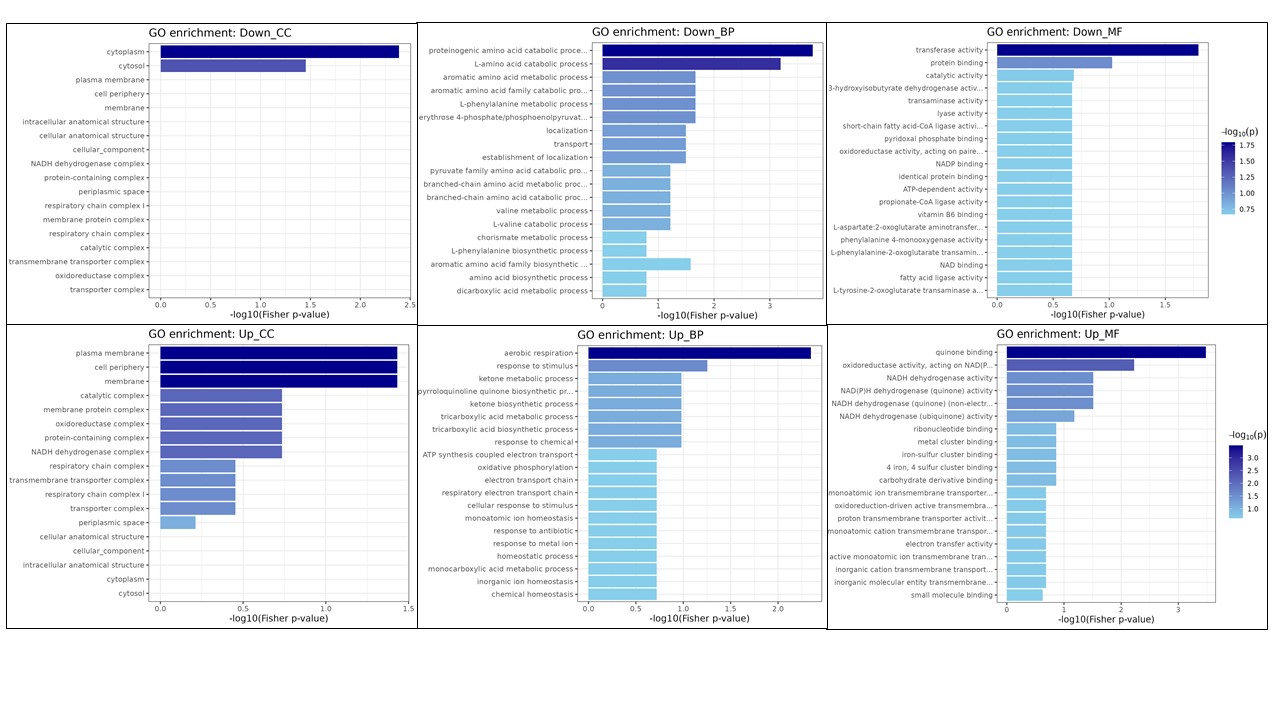

**Figure S7:** Function categorisation of DEGs in gene ontology (GO). The results were summarised in three main categories: biological processes, molecular function and cellular component.

**Table S4:** Overall Gene Ontology (GO) enrichment analysis of DEGs

| **Gene** | **log2FoldChange** | **Product** |
| --- | --- | --- |
|  | 3.229575835 | hypothetical protein |
|  | 3.177317772 | hypothetical protein |
|  | 2.842382553 | DUF485 domain-containing protein |
|  | 2.752890844 | Methyltransferase type 11 domain-containing protein |
|  | 3.548147201 | DUF1652 domain-containing protein |
|  | 2.65495853 | Methyltransferase type 11 domain-containing protein |
|  | -3.232069946 | FAD dependent oxidoreductase domain-containing protein |
| cioA | -2.823928315 | cyanide-insensitive cytochrome bd quinol oxidase subunit I |
|  | 3.382978121 | DUF1652 domain-containing protein |
| gntP | -4.206388693 | GntP family permease |
|  | 2.439890493 | histidine kinase |
|  | 2.528766638 | Lipoprotein |
| mmsA | -2.30655967 | Methylmalonate-semialdehyde dehydrogenase [acylating] |
|  | -3.158442944 | FAD dependent oxidoreductase domain-containing protein |
|  | 2.503081642 | L-lactate permease |
|  | 2.490488688 | Lipoprotein |
| mmsB | -2.369459924 | 3-hydroxyisobutyrate dehydrogenase |
| acs | 2.197816079 | acetate--CoA ligase |
| phhC | -2.219637142 | Aromatic-amino-acid aminotransferase |
|  | -2.222591619 | aldehyde dehydrogenase family protein |
|  | -2.719612173 | (S)-ureidoglycine aminohydrolase cupin domain-containing protein |
| phhC | -2.188958078 | Aromatic-amino-acid aminotransferase |
| mmsB | -2.323132123 | 3-hydroxyisobutyrate dehydrogenase |
|  | 2.461687852 | hybrid sensor histidine kinase/response regulator |
|  | 2.402729204 | hypothetical protein |
|  | -2.824516458 | (S)-ureidoglycine aminohydrolase cupin domain-containing protein |
| nuoM | 2.318420162 | NADH-quinone oxidoreductase subunit M |
|  | -2.337266902 | Pterin-4-alpha-carbinolamine dehydratase |
|  | -2.270904847 | Pterin-4-alpha-carbinolamine dehydratase |
|  | 2.560062371 | Sel1 domain-containing protein |
|  | 2.193172449 | Methyl-accepting chemotaxis protein |
|  | -2.456229189 | DUF3509 domain-containing protein |
|  | 2.474850089 | Sel1 domain-containing protein |
| phhA | -2.135008876 | phenylalanine 4-monooxygenase |
| phhA | -2.164976599 | phenylalanine 4-monooxygenase |
| lctP | 2.36040362 | lactate permease LctP family transporter |
| nuoN | 2.242174845 | NADH-quinone oxidoreductase subunit NuoN |
|  | 2.362741688 | UPF0337 protein PA4738 |
|  | 2.372601118 | UPF0337 protein PA4738 |
| lldD | 2.197971073 | FMN-dependent L-lactate dehydrogenase LldD |
|  | 2.481667868 | LTXXQ domain protein |
|  | -2.272373289 | Uncharacterized protein PA3568 |
|  | -2.332164588 | DUF3509 domain-containing protein |
| nuoH | 2.165652873 | NADH-quinone oxidoreductase subunit NuoH |
|  | 2.253052366 | hypothetical protein |
|  | -2.544635477 | DUF1654 domain-containing protein |
| nuoH | 2.246361379 | NADH-quinone oxidoreductase subunit NuoH |
| nuoL | 2.200482327 | NADH-quinone oxidoreductase subunit L |
| lldD | 2.240321274 | FMN-dependent L-lactate dehydrogenase LldD |
|  | -2.172025988 | Uncharacterized protein PA3568 |
| nuoJ | 2.190054075 | NADH-quinone oxidoreductase subunit J |
|  | -2.492358746 | DUF1654 domain-containing protein |
| carO | 2.062725217 | Calcium-regulated OB-fold protein CarO |
|  | 2.173045423 | BON domain-containing protein |
| carO | 2.039283809 | Calcium-regulated OB-fold protein CarO |
|  | 2.727747883 | Putative major facilitator superfamily transporter |
|  | 2.533665172 | DUF485 domain-containing protein |
| actP | 2.024957831 | Cation/acetate symporter ActP |
|  | 2.560316704 | DUF1682 domain-containing protein |
|  | 2.432539966 | hypothetical protein |
| gntP | -4.307592151 | GntP family permease |
| gcvT | -2.000589994 | glycine cleavage system aminomethyltransferase GcvT |
|  | -2.750221508 | Cyanide insensitive terminal oxidase, subunit III |
|  | 2.088457468 | Succinyl-CoA:coenzyme A transferase |
|  | 2.123757069 | BON domain-containing protein |
|  | -2.775371741 | Cyanide insensitive terminal oxidase, subunit III |
|  | 2.104293705 | NADH-quinone oxidoreductase |
|  | -2.498725671 | ABM domain-containing protein |
| piuC | 2.078381772 | Fe2+-dependent dioxygenase |
| nuoI | 2.184664564 | NADH-quinone oxidoreductase subunit NuoI |
| nuoI | 2.116557563 | NADH-quinone oxidoreductase subunit NuoI |
|  | 2.259547688 | putative membrane transporter protein |
| pqqE | 2.178576591 | pyrroloquinoline quinone biosynthesis protein PqqE |
|  | -2.362354819 | DUF2845 domain-containing protein |
| bdhA | -2.160857029 | 3-hydroxybutyrate dehydrogenase |
| pigA | 2.070528651 | biliverdin-producing heme oxygenase PigA |
|  | 2.216648572 | Yip1 domain-containing protein |
|  | 2.209275588 | NADH dehydrogenase I chain L |
|  | 2.12319044 | Cytochrome b561 bacterial/Ni-hydrogenase domain-containing protein |
|  | 2.070330444 | Yip1 domain-containing protein |
|  | -2.435679622 | Extracellular solute-binding protein |
| pqqB | 2.551939142 | pyrroloquinoline quinone biosynthesis protein PqqB |
|  | 2.004681725 | aminotransferase class V-fold PLP-dependent enzyme |
|  | 2.021106908 | putative membrane transporter protein |
|  | -2.064845603 | DUF1329 domain-containing protein |
| pqqC | 2.246736074 | pyrroloquinoline-quinone synthase PqqC |
|  | 2.085517012 | Cytochrome b561 bacterial/Ni-hydrogenase domain-containing protein |
|  | -2.348858731 | Extracellular solute-binding protein |
| nuoK | 2.125081695 | NADH-quinone oxidoreductase subunit NuoK |
|  | 2.123950806 | (Fe-S)-binding protein |
| mdcA | -2.668972877 | malonate decarboxylase subunit alpha |
|  | -2.483401769 | Secreted protein |
|  | -2.205771528 | Amino acid permease/ SLC12A domain-containing protein |
|  | 2.115156543 | UPF0391 membrane protein PLES_58781 |
|  | 2.069535753 | (Fe-S)-binding protein |
| nuoK | 2.093720265 | NADH-quinone oxidoreductase subunit NuoK |
|  | 2.056152426 | UPF0391 membrane protein PLES_58781 |
|  | -2.166030672 | Re/Si-specific NAD(P)(+) transhydrogenase subunit alpha |
|  | 2.553282834 | Putative major facilitator superfamily transporter |
| pqqD | 2.3355754 | pyrroloquinoline quinone biosynthesis peptide chaperone PqqD |
|  | -4.441047756 | hypothetical protein |
|  | -2.06589559 | Re/Si-specific NAD(P)(+) transhydrogenase subunit alpha |
|  | 2.592307981 | Secreted protein |
| yohK | 2.137761492 | Inner membrane protein YohK |
| mdcA | -2.496345084 | malonate decarboxylase subunit alpha |
|  | -2.147205715 | Secreted protein |
| flp | -2.159602056 | type 4b pilus Flp major pilin |
|  | 2.378207606 | TonB-dependent siderophore receptor |
|  | -2.512516987 | putative hydro-lyase |
| flp | -2.123626412 | type 4b pilus Flp major pilin |
|  | -2.434126047 | DUF1329 domain-containing protein |
| piuC | 2.123142408 | Iron-uptake factor PiuC |
|  | -2.172422877 | putative hydro-lyase |
|  | 2.09936678 | acetyl-CoA hydrolase/transferase C-terminal domain-containing protein |
|  | 2.167141802 | hypothetical protein |
|  | 2.365949532 | Secreted protein |
|  | 2.91183154 | Coenzyme PQQ synthesis protein B |
| ureE | -2.237655266 | urease accessory protein UreE |
| araC | -2.029586981 | AraC family transcriptional regulator |
| fabG | -2.372567783 | 3-oxoacyl-[acyl-carrier-protein] reductase FabG |
|  | -2.513382868 | L-serine ammonia-lyase, iron-sulfur-dependent, subunit alpha |
|  | -2.105497456 | Aromatic amino acid permease |
|  | -2.265568802 | CoA transferase |
|  | -2.359838121 | hypothetical protein |
| cyoA | -2.618184249 | ubiquinol oxidase subunit II |
|  | -2.169061195 | MFS transporter |
|  | -2.18977517 | hypothetical protein |
|  | -2.045798938 | Amino acid permease/ SLC12A domain-containing protein |
| ureF | -2.330865483 | urease accessory protein UreF |
|  | -2.400087754 | hypothetical protein |
|  | -2.787148165 | hypothetical protein |
|  | -2.24794783 | hypothetical protein |
|  | -2.459281258 | DUF2790 domain-containing protein |
|  | 2.485744349 | TonB-dependent siderophore receptor |
